# Microspatial partitioning of insect-specific viromes and dengue virus transmission risk by *Aedes aegypt*i in Puerto Rico

**DOI:** 10.64898/2026.07.15.738651

**Authors:** Bright Agbodzi, Luis Alonso-Palomares, Henry J. Barton, Nicole Nazario-Maldonado, Joanelis Medina Quintana, Nexilianne Borrero-Segarra, John F. Williams, Vanessa Rivera-Amill, Robert Rodriguez-Gonzalez, Grayson Brown, Rhoel R. Dinglasan

## Abstract

Mosquito-borne arboviruses remain a major global health threat. Although insect-specific viruses (ISVs) may influence arbovirus transmission across geographic regions, their diversity within *Aedes aegypti* populations at microspatial scales remains poorly understood. We hypothesized that local environmental variation shapes the *A. aegypti* core virome, producing distinct ISV profiles in rural and urban populations in Puerto Rico. Metatranscriptomic RNA sequencing of F0 mosquitoes identified thirteen ISVs with habitat-specific virome composition. Partitiviridae was the dominant viral family, while Humaita-Tubiacanga virus (HTV) and Phasi Charoen-like phasivirus (PCLV) exhibited the highest abundance, persistence, and prevalence in urban laboratory colonies. Urban *A. aegypti* also showed significantly higher dengue virus-1 (DENV-1) infection rates than rural populations, although saliva positivity remained low and did not differ significantly between the two groups. These findings indicate that microspatial ecological variation shapes the *A. aegypti* virome, but the association of HTV/PCLV with DENV-1 transmission risk is likely multifactorial and nuanced.

## INTRODUCTION

Mosquito-borne diseases (MBDs) are a major public health burden, affecting populations across tropical, subtropical, and temperate regions. The World Health Organization (WHO) estimates that about half of the world’s population is at risk of dengue fever, which is caused by dengue viruses (DENV), subtypes 1-4, a flavivirus primarily transmitted by the *Aedes aegypti* mosquito, causing 96 million symptomatic cases and 40,000 deaths every year^1,2^. Fully understanding the biology of the vector is critical for providing insights that could inform the development of novel biocontrol strategies against MBDs. While the scientific community has examined the innate immune response of mosquito vectors to understand vector host-arbovirus interactions, over the past decade appreciation of the influence of a mosquito’s microbiome (including bacteria, parasites, and viruses) on vector competence (the intrinsic ability of an arthropod to acquire and and transmit a pathogen)^3^ has grown. By understanding the basics of vector host-microbiome interactions, we can potentially take advantage of this relationship to develop new vector control interventions to prevent MBD transmission^4–6^. Next-generation sequencing (NGS) has enabled the discovery of numerous insect-specific viruses (ISVs), traditionally considered restricted to insects^4,7–9^; however, some, including the Guadeloupe mosquito virus (GMV), have recently been detected in humans^10^. Some of these ISVs have been shown to modulate vector competence by enhancing or decreasing the replication success of medically important arboviruses^6,7,11–14^. However, the variation in the distribution and composition of ISVs in natural mosquito populations remains poorly characterized, as much of the mosquito metagenomic microbiome data has been limited to established laboratory colonies. Moreover, natural acquisition of ISVs by mosquitoes has been associated with transovarial (vertical) transmission from mother to offspring, and venereal transmission (from male to female)^8,15,16^; however, the efficiency of this process for different ISVs is uncertain.

How mosquito ecology, spatial partitioning within an endemic disease focus, habitat characteristics, and interaction with other insect species contribute to changes in ISV composition in a disease-endemic setting such as Puerto Rico is unclear.^17–19^ Do natural populations of *Aedes aegypti* exhibit distinct ISV profiles compared to long-term laboratory colonies? Could these differences underlie variations in their vector competence for different DENV serotypes? We investigated the ecological profile of ISVs in natural populations of *A. aegypti* from Puerto Rico, a Caribbean Island, with historical dengue endemicity since 1915 and cyclical, multi-serotype epidemics (DENV-1 to -4)^17–19^, often stemming from an initial outbreak in San Juan, Puerto Rico’s capital. A previous study that used RT-qPCR screening for ISVs found that several populations of *A. aegypti* from the main island, including from Culebra, an offshore island of Puerto Rico, were positive for Cell-Fusing Agent Virus (CFAV), the first flavivirus ISV recorded in natural mosquito populations ^20^. However, the influence of CFAV in vector competence remains equivocal ^21,22^, suggesting that different ISV profile might potentially influence recurrent DENV transmission in Puerto Rico. To date, no other studies have explored or captured the ISV profiles for mosquito populations in Puerto Rico’s main island. Herein, we used metatranscriptomics sequencing and functional studies to test the hypothesis that micro-spatial partitioning shapes virome diversity of natural *A. aegypti* populations, which in turn, directly influence vector competence and DENV transmission risk.

## METHODS

### Sampling

To select urban vs. rural habitat, we defined our study sites based on their 1) demographics (high vs. low population density) and geographical landscape (highly urbanized vs. low urbanized)^23^, 2) dengue incidence (2010-2020 reported local cases)^24^, and 3) availability of *A. aegypti* surveillance data and ecological studies^25^ (**Figure 1**). For urban habitats, we selected the communities of Reparto Metropolitano (R) and Cantera (C) in San Juan municipality (> 326,000 inhabitants), both highly urbanized communities, with historically high dengue incidence^26^ and an established mosquito surveillance system by the Puerto Rico Vector Control Unit (PRVCU)^25^. For rural habitats, we selected the communities of Providencia (P) and Lamboglia (L) in Patillas municipality (> 15,000 inhabitants) based on their low population density, and sporadic dengue cases. ArcGIS Pro was used to randomly select 10 houses for site visits in July 2021. However, due to logistical challenges in obtaining repeat resident authorization for collections, we reduced the number of sampling sites to 5 houses/site in March 2022,. Adult mosquitoes were collected using CDC light traps, while mosquito eggs were collected using ovitraps and used to establish field-derived colonies under laboratory conditions. Sampling was conducted in July 2021 and March 2022, and a subset of 12 adult *A. aegypti* females from each site/period (**Figure 1**) were selected for abdomen dissection to capture midgut ISVs. Sampling for vector competence and ISV prevalence work was done in March-April 2024.

**Figure 1.**
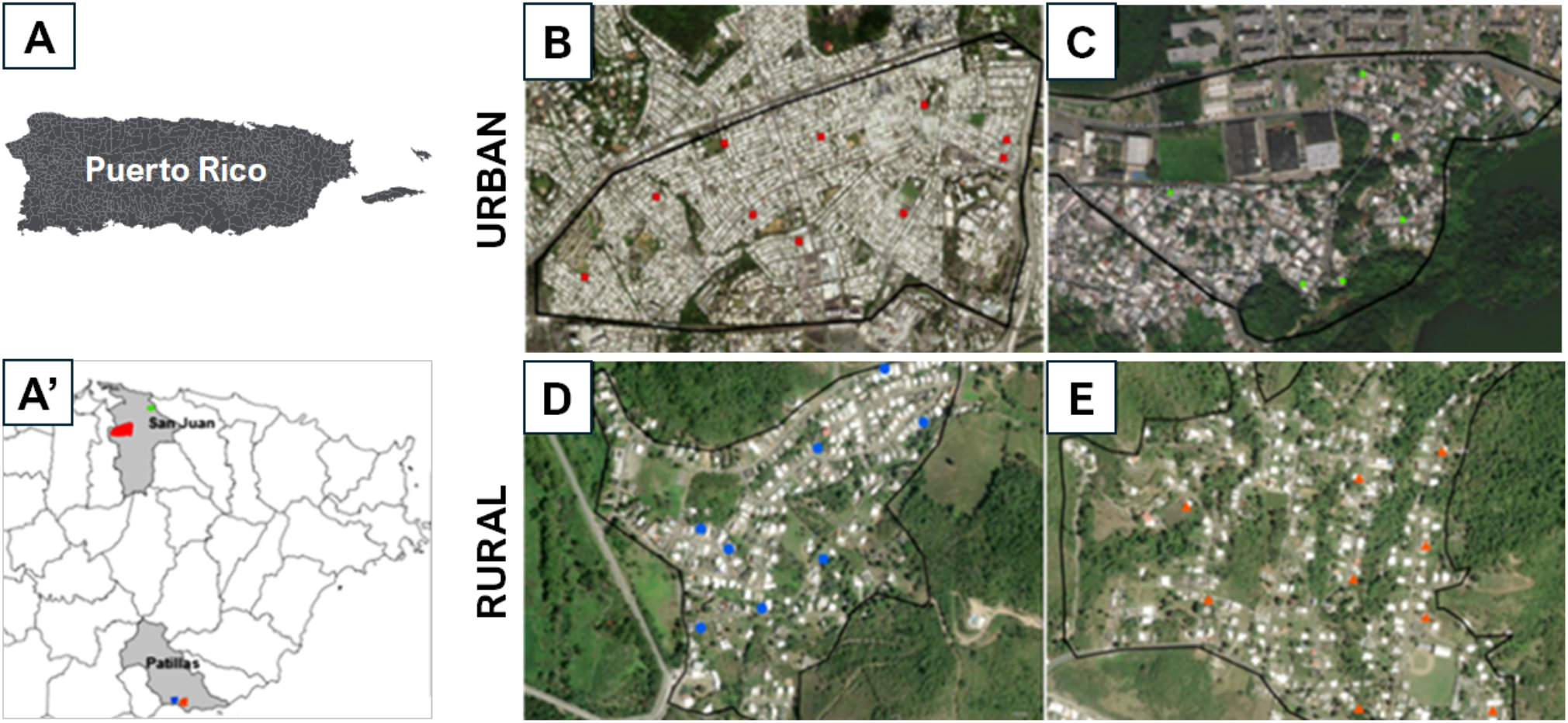
*Aedes aegypti* sampling sites in the eastern section of the main island of Puerto Rico. (A) Puerto Rico island group composed of the main large island and two smaller islands to the east (Culebra and Vieques). (A’) Municipalities of San Juan (Capital) and Patillas were targeted for sampling of *A. aegypti* mosquitoes from urban and rural sites. Sampling of *A. aegypti* larvae from individual homes in Reparto Metropolitano (B) and Cantera (C) in the municipality of San Juan, as well as Providencia (D) and Lamboglia (E) in the municipality of Patillas.

### Mosquito Maintenance

Mosquito colonies from rural and urban communities were established in four separate growth chambers in an Arthropod Containment Level 2 (ACL-2) facility. Mosquito eggs were hatched in deionized water (DI), and larvae were grown at a density of about 200 larvae per tray. Adult mosquitoes were morphologically identified to species using standard taxonomic keys. Adult mosquitoes were maintained on 10% sucrose solution while housed in 28 °C growth chambers at 80% relative humidity (RH) and a 12/12 light/dark cycle at the University of Florida as per standard procedures^27^. To maintain the colonies, females were blood-fed using a 1:1 mixture of O+ human red blood cells (RBCs) (Lifesouth Community Blood Centers, Gainesville, FL, USA) and heat-inactivated human serum (HHIS) delivered via a water-jacketed artificial membrane feeder maintained at 38 °C.

### ISV Prevalence and Generational Maintenance Testing

To assess the prevalence of ISVs in urban and rural mosquito colonies, we used the sequences of the four ISVs obtained from our metatranscriptomic analysis (Supplementary Table 1) to design the primers (Supplementary Table 3). The targeted viruses included *Aedes aegypti* toti-like virus (ATLV), Guadeloupe mosquito virus (GMV), Humaita-Tubiacanga virus (HTV) and Phasi Charoen-like virus (PCLV). PCR amplicons were validated by cloning using the TOPO™ TA Cloning Kit (Invitrogen, Thermo Fisher Scientific) following the manufacturer’s instructions. The insert identity was confirmed by Sanger sequencing. For prevalence screening, abdomens from 25 females and 25 males individual mosquitoes from both the F₀ and F₁ generations were tested for PCLV and HTV. In contrast, GMV and ATLV screening was performed using only F₀ mosquitoes. Abdomens were dissected and homogenized in 500 µL of TRIzol containing approximately 0.15 g of 0.5 mm glass beads (Next Advance, Cat. No. GB05) using a Bullet Blender® (Cat. No. BT24M) at speed 8 for 5 minutes. Total RNA was extracted using TRIzol™ Reagent (Thermo Fisher Scientific, Waltham, MA, USA) according to the manufacturer’s instructions.

### Virus Strain

The DENV-1 strain Haiti/1207/2014: KT279761.2^28^, which was isolated from a child in Haiti, was used in this study. A working virus stock was generated from two passages of the isolate in Vero E6 cells followed by 3 passages in C6/36 cells. Cell infection was monitored by observing cytopathic effect (CPE). Viral stocks were collected between 7- and 9-days post-inoculation when approximately 50% of the cells showed virus-specific CPE. The viral stock was tittered, stabilized with 10% *w*/*v* molecular grade trehalose, and cryopreserved as previously described^27^.

### Mosquito Infection and Tissue Collection

Lamboglia and Reparto colonies were selected to represent the urban and rural populations respectively. We used F₆ female mosquitoes for all experimental replicates. We included the laboratory strain A. *aegypti* ORL-D for comparison. For each replicate, 60 female mosquitoes were offered an infectious blood meal consisting of a 2:2:1 mixture of O+ human RBCs, DENV-1 (1 × 10¹⁰ PFU/mL), and HHIS to deliver an infectious dose of 4 × 10⁹ PFU/mL. The experiment was conducted in an Arthropod Containment Level 3 (ACL-3) facility. Fully engorged females were maintained on 10% sucrose at 28 °C and 80% relative humidity until 14 days post-infection (dpi). At 14 dpi, mosquitoes were deprived of sucrose for 6 hours prior to processing, then cold-anesthetized at 4 °C for 10 minutes and kept on ice. Saliva was collected by first removing the legs and wings, followed by inserting the proboscis into a capillary tube containing 10 µL of a solution composed of 1 part human RBCs and 3 parts HHIS for 30 minutes^27^. Following saliva collection, abdomens were dissected and transferred into 1.5 mL safe-lock microcentrifuge tubes (Eppendorf, Cat. No. 0030123328) containing 500 µL of Dulbecco’s Modified Eagle Medium (DMEM; Gibco, Thermo Fisher Scientific, Waltham, MA, USA). supplemented with 3% fetal bovine serum (FBS; Sigma-Aldrich, St. Louis, MO, USA)., 1× penicillin/streptomycin, and approximately 0.15 g of 0.5 mm glass beads (Next Advance, Cat. No. GB05). Samples were stored at −80 °C until further processing. When more than 30 mosquitoes survived at 14 dpi, a maximum of 30 individuals per group were processed for downstream analyses.

Infection rate (IR) was determined as the proportion of mosquitoes with DENV-1-positive abdomen among tested mosquitoes. Transmission potential (TP) represents the proportion of mosquitoes with DENV-1-positive saliva among mosquitoes with abdominal infection.

### RNA Extraction and RT-qPCR

The samples were thawed on ice and homogenized in a Bullet Blender^®^ at speed 8 for 5 min. The samples were then centrifuged at 3800×*g* for 3 min at 4 °C to pellet debris. RNA was extracted from the virions in the supernatant using a QIAmp Viral RNA Mini Kit (Qiagen, Cat. 52906), following the manufacturer’s instructions. Primer information for DENV-1 testing is available in Supplementary Table 3. One-step quantitative reverse transcription polymerase chain reaction (RT-qPCR) was performed using the UltraPlex 1-Step ToughMix (Quantabio, Beverly, MA, USA). The thermocycling conditions consisted of an initial reverse transcription step at 50 °C for 10 min, followed by an initial denaturation at 95 °C for 2 min, and 45 amplification cycles of 95 °C for 15 s and 60 °C for 45 s. Samples with a quantification cycle (Cq) value ≤ 38 were considered positive. Amplification controls (synthetic viral nucleic acids) were included on each plate from two biological replicates, with duplicate replicates per sample. Samples we run in duplicate using the Azure Cielo Real-Time PCR system (Azure Biosystems Mod. AIQ060).

### Metatranscriptomic Sequencing

Total RNA was extracted from the abdomen of female mosquitoes using the Qiagen AllPrep DNA/RNA mini kit, following manufacturer’s instructions. A pool of 3-6 samples was made from individuals with RNA eluates (RNA Concentration > 50 ng/µl) from each site/period. Libraries were constructed (Novogene USA). We used our established RNASeq metagenomic assembly pipeline^15^ to further investigate the mosquito virome of *A. aegypti* adult females (collected as F_0_ larvae) from the two urban communities in San Juan and two rural communities in Patillas, Puerto Rico. For comparison, we included two *Aedes aegypti* Orlando laboratory strains: ORL-D from the DeGennaro Lab at Florida International University and the Orlando strain from the Center for Medical, Agricultural, and Veterinary Entomology (CMAVE) at the University of Florida.

### Bioinformatic Analysis

#### Read preparation

Raw paired-end Illumina sequencing reads were processed using an integrated bioinformatic workflow comprising quality trimming, host read removal, de novo assembly, viral annotation, and quantitative read mapping. Adapter sequences and low-quality bases were removed using BBDuk (BBTools suite), applying right-end quality trimming (Q ≥ 20), minimum read length filtering (≥ 50 bp), and adapter reference matching.

Trimmed reads were mapped to the *Aedes aegypti* Liverpool reference genome (AaegL5.1) using BBMap^29^. Host-aligned reads were discarded, and unmapped paired reads were retained for downstream metagenomic analyses.

#### Alignment and annotation

Unmapped reads (non-host) were assembled *de novo* using SPAdes version 3.15.5^30^ with the --meta flag specified. The assembled contigs were filtered to retain sequences ≥ 300 bp. Protein-level similarity searches were performed against the NCBI non-redundant (NR) protein database using DIAMOND blastx^31^, retaining up to 10 target hits per query with an E-value threshold of 1×10^-10^ and minimum amino acid identity of 95%. DIAMOND outputs were generated in both DAA format for downstream taxonomic classification and tabular format (outfmt 6) for audit and downstream parsing.

#### Lowest common ancestor (LCA) analysis

Taxonomic classification was performed using MEGAN v6 ^32^ via a lowest common ancestor (LCA) algorithm applied to DIAMOND DAA outputs. Viral contigs were identified by filtering assignments to the viral taxonomic lineage. Viral contigs were exported on a per-sample basis for quantitative analysis. Contigs were retained as high-confidence viral candidates if their taxonomic lineage contained the term “Viruses,” supported by ≥ 20 mapped reads and a breadth of coverage ≥ 0.90. Viral relative abundance was calculated as the number of reads mapping to viral contigs divided by the total number of reads mapping to all passing viral contigs within each sample. The relative abundances of the passing viruses were then visualized.

#### Phylogenetic analysis

Draft genome segments were used to query the NCBI blast database to select representative sequences for comparison. Sequences were aligned using MUSCLE, implemented in MEGA X. To correct for the effects of ambiguous alignments due to the polymorphisms in 5’ and 3’ untranslated regions, the sequences were trimmed to the open reading frames (ORFs), and all subsequent phylogenetic analyses were conducted on the ORFs. A maximum likelihood phylogenetic analysis was conducted using IQ-TREE with simultaneous best-fit model selection and ultrafast bootstrapping (N = 1,000).

### Histone H4 analysis

Thirty female mosquitoes from the Lamboglia (rural) and Reparto (urban) colonies were offered an infectious DENV-1 blood meal as described above. Uninfected controls were treated similarly, except that the virus stock was replaced with DMEM. At 14 dpi, mosquito abdomens were dissected on ice and stored at −80 °C until RNA extraction. Abdomens were homogenized in 500 µL of TRIzol™ Reagent containing approximately 0.15 g of 0.5 mm glass beads using a Bullet Blender® at speed 8 for 5 min. Total RNA was extracted using TRIzol Reagent according to the manufacturer’s instructions. DENV-1 infection was confirmed with RT-qPCR prior to downstream analysis. For each colony, RNA from eight infected individual mosquitoes was pooled by concentration for *histone H4* gene expression analysis. Total RNA from pools were reverse transcribed into first-strand cDNA using the SuperScript IV First-Strand Synthesis System (Thermo Fisher Scientific, Waltham, MA, USA) according to the manufacturer’s instructions with random hexamer priming. Synthesized cDNA was quantified using a NanoDrop Spectrophotometer (Thermo Fisher Scientific, Waltham, MA, USA), diluted as needed. *Histone H4* gene expression was quantified in infected and uninfected mosquitoes from the Lamboglia (rural) and Reparto (urban) colonies by RT-qPCR using the Azure Cielo Real-Time PCR System (Azure Biosystems, Dublin, CA, USA). Primers for *histone H4* amplification were previously described^6^, and the ribosomal protein gene *rpl32* was used as the endogenous control. Relative gene expression was calculated using the comparative 2-ΔΔCt method, with uninfected mosquitoes serving as the calibrator group. Three independent biological replicates were analyzed per treatment group.

### Statistical Analysis

Vector competence outcomes—specifically infection status (infected = 1, non-infected = 0) and transmission potential (saliva positivity: positive = 1, negative = 0)—were modeled separately using multilevel logistic regression with a binomial error distribution and logit link function (lme4:glmer). Mosquito strain (ORL-D, Reparto, and Lamboglia) was included as a fixed effect, with Reparto (urban strain) serving as the reference group to assess differences against ORL-D (laboratory strain) and Lamboglia (rural strain), while experimental replicate was included as a random intercept to account for batch effects. Odds ratios (ORs) and 95% confidence intervals (CIs) were calculated from model coefficients, and pairwise comparisons among strains were performed using estimated marginal means with Tukey-adjusted *p*-values. Viral genome copy numbers were log₁₀ transformed and analyzed using non-parametric tests due to non-normal distribution; differences in viral load among locations were assessed for each combination of sex, generation, and virus type using the Kruskal-Wallis test (restricting analysis to location groups with sample sizes > 9), followed by pairwise Wilcoxon rank-sum tests with Benjamini-Hochberg correction for significant overall differences (*p* < 0.05). ISV prevalence was estimated as percentages with exact 95% CIs calculated using the binom.test function. Finally, for *histone H4* analysis, statistical comparisons between infected and uninfected mosquitoes within each colony were performed on ΔCt values using paired Student’s *t*-tests, with *p*-values adjusted for multiple comparisons using the Benjamini–Hochberg method. All statistical analyses were performed in R (version 4.5.2) with significance defined at α = 0.05 and visualizations were generated using the ggplot2, ggh4x, ggpubr, and rstatix packages.

## RESULTS

### Microbial read distribution varies between rural and urban *A. aegypti*

The proportion of filtered non-host reads ranged from 6% to 22% of total raw reads (Supplementary Table 1). Among non-host reads, the proportion aligning to the reference database varied from 2.7% to 55% across samples (Supplementary Table 1). Of the database-aligned reads, viral sequences predominated in urban samples, accounting for 72–92% of aligned reads, whereas rural samples contained substantially lower viral proportions (12–40%). Notably, the two laboratory strains exhibited markedly different viral contents, with viral reads representing 38.8% and 0.06% of database-aligned reads, respectively (Supplementary Table 1). In contrast, bacterial and eukaryotic sequences constitute a greater proportion of database-aligned reads in rural than in urban samples (Supplementary Table 1). Viral assembly reads metrics are presented in (Supplementary Table 2).

### A. aegypti-associated ISV profiles

In total, we detected 13 viruses in this study, which partitioned between rural and urban sampling sites on the main island of Puerto Rico (**Table 1**, **Figure 2**). Of these, the most represented families were *Partitiviridae* and *Totiviridae* (2 viruses each). Other virus families include *Flaviviridae, Phenuiviridae, Orthomyxoviridae, Rhabdoviridae, Virgaviridae and Picobirnaviridae.* Four of the detected viruses remain unclassified. Overall, both Phasi Charoen-like phasivirus (PCLV; *Phenuiviridae*) and Humaita-Tubiacanga virus (HTV; unclassified virus) had the highest number of virus sequences. The cell-fusing agent virus (CFAV) was the only virus detected across all mosquito populations/strains in our study.

**Figure 2.**
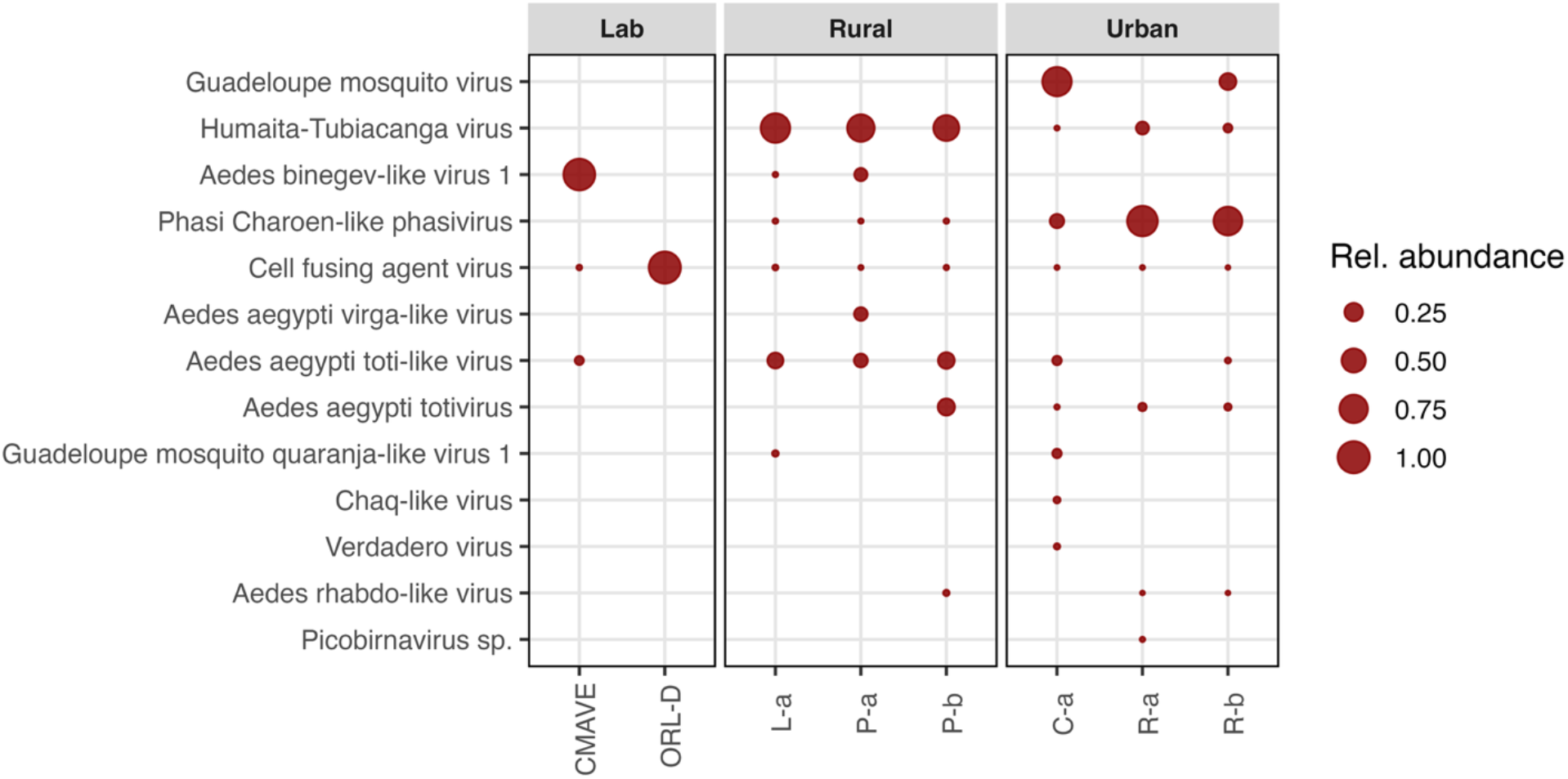
Relative abundance of aligned viral reads (per virus species) from field *A. aegypti* female abdomens from rural (Providencia: sub-petitioned into P-a and P-b & Lamboglia: L-a) and urban (Reparto Metropolitano: sub-petitioned into R-a and R-b & Cantera: C-a) sites in Puerto Rico. Two laboratory strains, ORL-D and CMAVE were added for context. The y-axis is ordered alphabetically by taxon, while the x-axis represents sampling location. Relative read abundance per sample is depicted by the size of circle as indicated by the abundance key.

**Table 1.** Classification of insect-specific viruses that were identified in field-collected *Aedes aegypti*.

| Virus | Classification | Strand (polarity) | Rural/Urban | Notable field description<br>(Country; citation) |
| --- | --- | --- | --- | --- |
| Verdadero virus | *Partitiviridae* | dsRNA | Urban | Mexico <sup>53</sup> |
| *Aedes binegev*-like virus 1 | unclassified | ssRNA (+) | Both | Unknown<sup>41</sup> |
| Phasi Charoen-like virus | *Phenuiviridae* | ssRNA (-) | Both | Thailand <sup>54</sup> |
| *Aedes aegypti* totivirus | *Totiviridae* | dsRNA | Both | Ghana<sup>55</sup> |
| Humaita-Tubiacanga virus | unclassified | ssRNA (+) | Both | Brazil <sup>35</sup> |
| Guadeloupe mosquito virus | unclassified | ssRNA (+) | Urban | Guadeloupe <sup>36</sup> |
| Guadeloupe mosquito quaranja-like virus 1 | *Orthomyxoviridae* | ssRNA (-) | Both | Guadeloupe <sup>36</sup> |
| Chaq-like virus | *Partitiviridae* | dsRNA | Urban | Nigeria<sup>44</sup> |
| Cell fusing agent virus | *Flaviviridae* | ssRNA (+) | Both | Puerto Rico <sup>20</sup> |
| *Aedes rhabdo*-like virus | *Rhabdoviridae* | ssRNA (-) | Rural | Brazil (Unpublished) |
| Picobirnavirus spp. | *Picobirnaviridae* | dsRNA | Urban | - |
| *Aedes aegypti* toti-like virus | *Totiviridae* | dsRNA | Both | Brazil <sup>56</sup> |
| *Aedes aegypti* virga-like virus | *Virgaviridae* | ssRNA (+) | Rural | Ghana<sup>55</sup> |

Dashes (-) indicate data is unavailable.

### Virome composition differs between rural and urban *A. aegypti*

Of the 13 viruses detected, eight were common to both rural and urban sites; however, four were present only in urban *A. aegypti* and absent from the rural samples (**Figure 2**). These viruses include Verdadero virus, Picobirnavirus spp., GMV, and Chaq-like virus. Conversely, two viruses (*Aedes* binegev-like virus 1 and Croada virus) were detected in the rural samples but were absent from the urban sites. The laboratory strains exhibited distinct virome compositions: ORL-D harbored only CFAV, whereas CMAVE contained multiple viruses, including CFAV, *Aedes* binegev-like virus 1, and unclassified viral reads. Among the rural *A. aegypti* collected, HTV was the most predominant virus (>60%); however, PCLV was detectable at very low levels (<1% reads). In contrast, for urban *A. aegypti*, PCLV was the most predominant virus, with >75% of reads (in two of three pools), while HTV reads were low overall (<8%).

### Phylogenetic analysis

The phasivirus PCLV is a tri-segmented, negative-sense RNA virus in the order *Bunyavirales*^33,34^. Phylogenetic analysis of the three PCLV segments (**Figure 3**) showed a similar clustering pattern, with strains from the current study clustering closely together. The closest GenBank matches for the L segments were laboratory strains from the United States of America (USA), Australia, and the United Kingdom (MH310079.1, MH237599.1 and KU936057.1, respectively) with nucleotide identities of ∼98.0%. The closest wild sequences were the 2017 strains from Guadeloupe. The segment M sequences were closest to the NCBI reference sequence NC_038261.1 and to MN692604, with nucleotide percent identities of 97.6 and 97.5, respectively. These two sequences originate from Brazil. For the S segment, the closest matches were Asian strains with a nucleotide identity of ∼ 97%.

**Figure 3.**
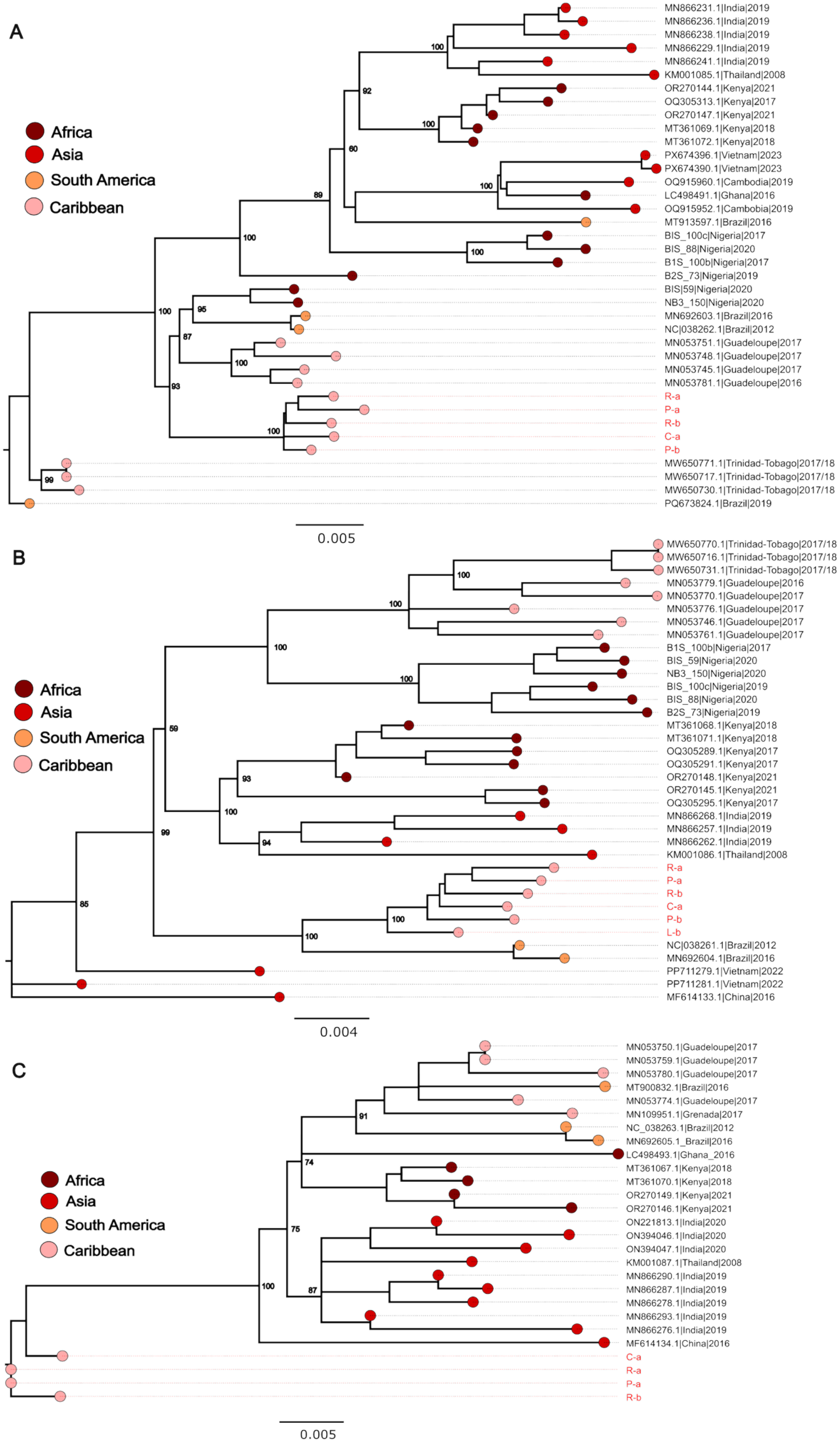
Phasi Charoen-like virus (PCLV) phylogeny. PCLV L Segment (RdRp) (A), Segment (GP gene) (B) and S segment (C). Tree model inference and phylogeny were simultaneously conducted in IQ-TREE v1.6.1, executing 1000 bootstrap replicates. The geographical origins of sequences are color coded and indicated in the Key. Critical nodes are labelled with bootstrap values. The tree was visualized in FigTree v1.4.4.

Although currently unclassified, HTVs, which were originally isolated from *A. aegypti* from Brazil, are positive-sense (+), single-stranded (ssRNA) bi-segmented viruses ^35^. The phylogeny of the sampled HTV revealed that for both segments 1 and 2, our sequences formed a separate cluster (**Figure 4**). For segment 1 (RNA-dependent RNA polymerase, RdRp), the closest match was a 2012 strain from Brazil (KR003801.1), with a nucleotide identity of ∼98.5%. Segment 2 (Capsid) sequences were the closest to 2016/17 Guadeloupe strains, with nucleotide identity of ∼98%.

**Figure 4.**
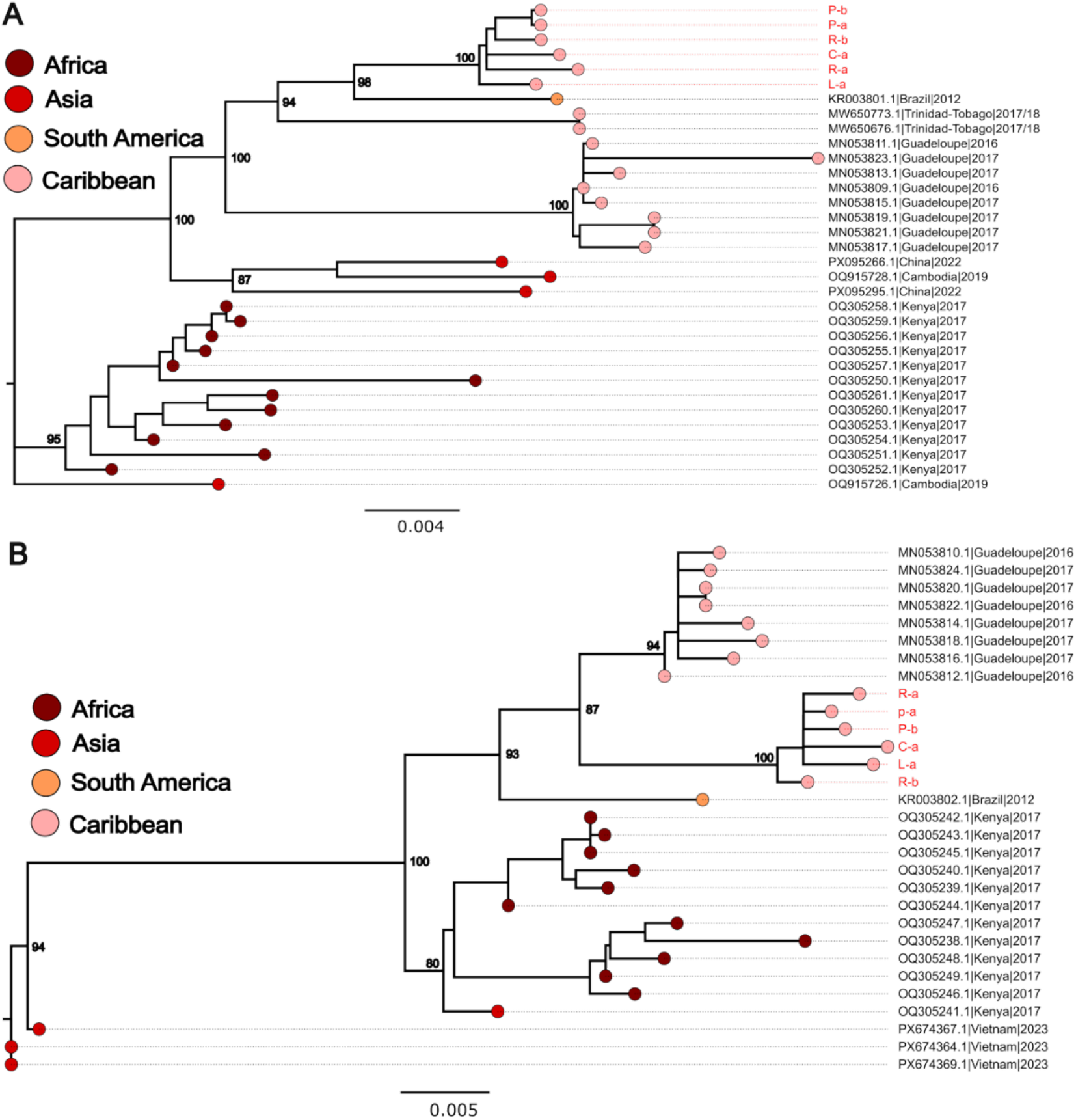
Humaita-Tubiacanga (HTV) phylogeny. HTV Segment 1 (RdRp) (A), Segment 2 (Capsid) (B). Tree model inference and phylogeny were simultaneously conducted in IQ-TREE v1.6.1, executing 1000 bootstrap replicates. The geographical origins of sequences are color coded and indicated in the Key. Critical nodes are labelled with bootstrap values. The tree was visualized in FigTree.

GMV is an unclassified bi-segmented, sobemo-like positive-sense RNA virus that groups with *Solemoviridae*-related viruses ^36^. The GMV phylogeny for segment 1 (**Figure 5**) revealed that our sequences did not cluster together, nor with other reference strains; however, our sequences shared ∼99% nucleotide identity with 2017/18 strains from the USA. Segment 2 sequences clustered with 2017/18 US strains (nucleotide identity of ∼99%).

**Figure 5.**
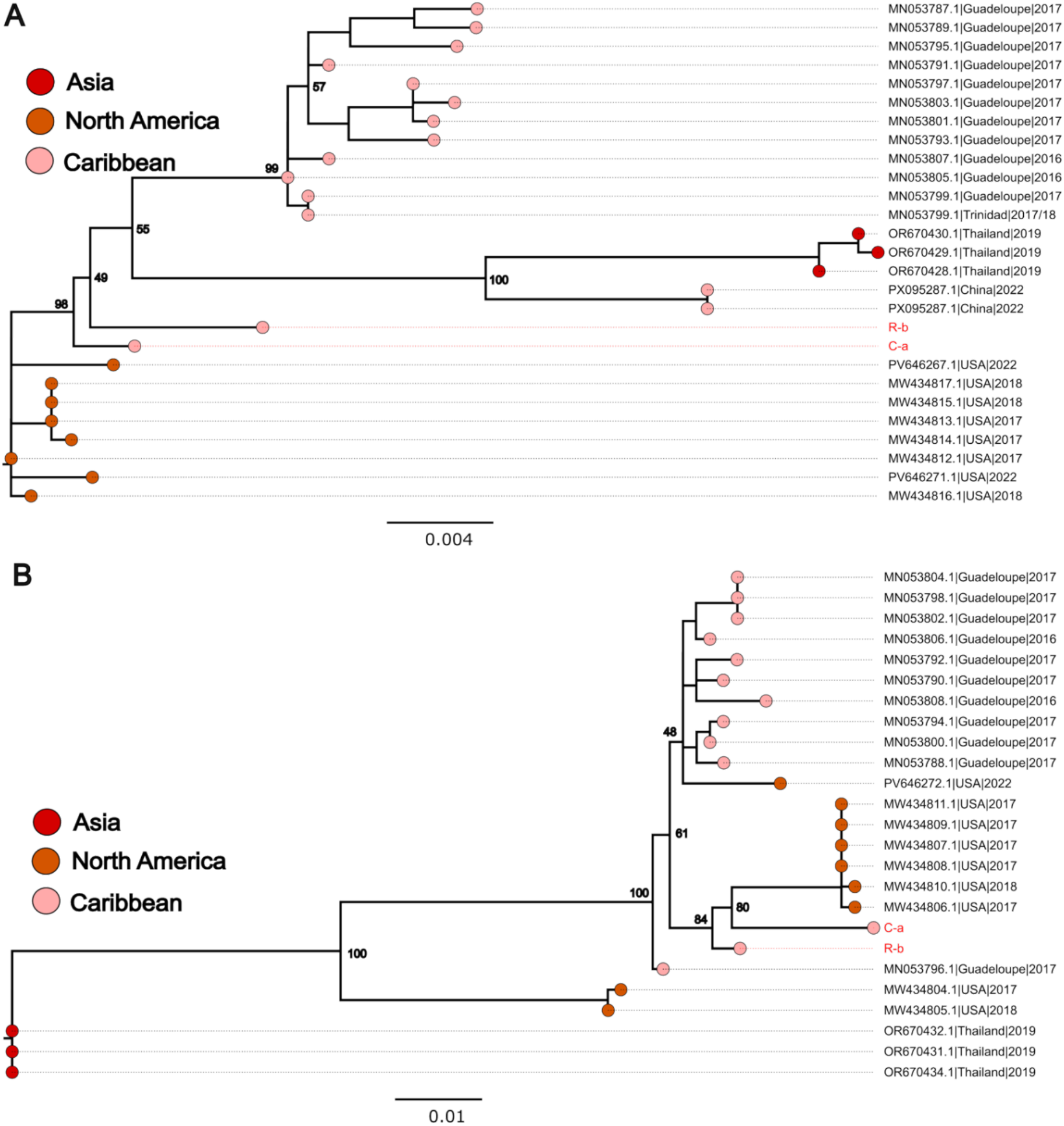
Guadeloupe mosquito virus (GMV) phylogeny. GMV Segment 1 (RdRp) (A), Segment 2 (Capsid) (B). Tree model inference and phylogeny were simultaneously conducted in IQ-TREE v1.6.1, executing 1000 bootstrap replicates. The geographical origins of sequences are color coded and indicated in the Key. Critical nodes are labelled with bootstrap values. The tree was visualized in FigTree.

*Aedes aegypti* toti-like virus (ATLV) is an insect-associated, double-stranded RNA virus belonging to the family *Totiviridae*, characterized by a non-segmented genome encoding a capsid protein and an RNA-dependent RNA polymerase^37^. The ATLV strains identified in this study shared ∼98% nucleotide identity with 2017 strains from Guadeloupe. Phylogeny was not explored because of a lack of available reference strains in GenBank as of April 2026.

### ISV Prevalence and Virus Load Vary Between Urban and Rural Populations

In urban-derived colonies (Cantera and Reparto), PCLV prevalence was consistently high across two generations (F_0_ and F_1_) and sexes, ranging from 96–100%. Similarly, HTV infection prevalence in urban samples was high, between 84–100%. In contrast, rural-derived colonies (Providencia and Lamboglia) exhibited greater variability in infection prevalence (**Figure 6A**). We observed that the infection prevalence of PCLV ranged from 44% to 100%, with the lowest observed in Lamboglia males (44–48%) compared to Providencia (84–100%). HTV prevalence in rural-derived populations was more heterogeneous, ranging from as low as 40% in Providencia F_0_ males to 96% in F_1_ females.

**Figure 6.**
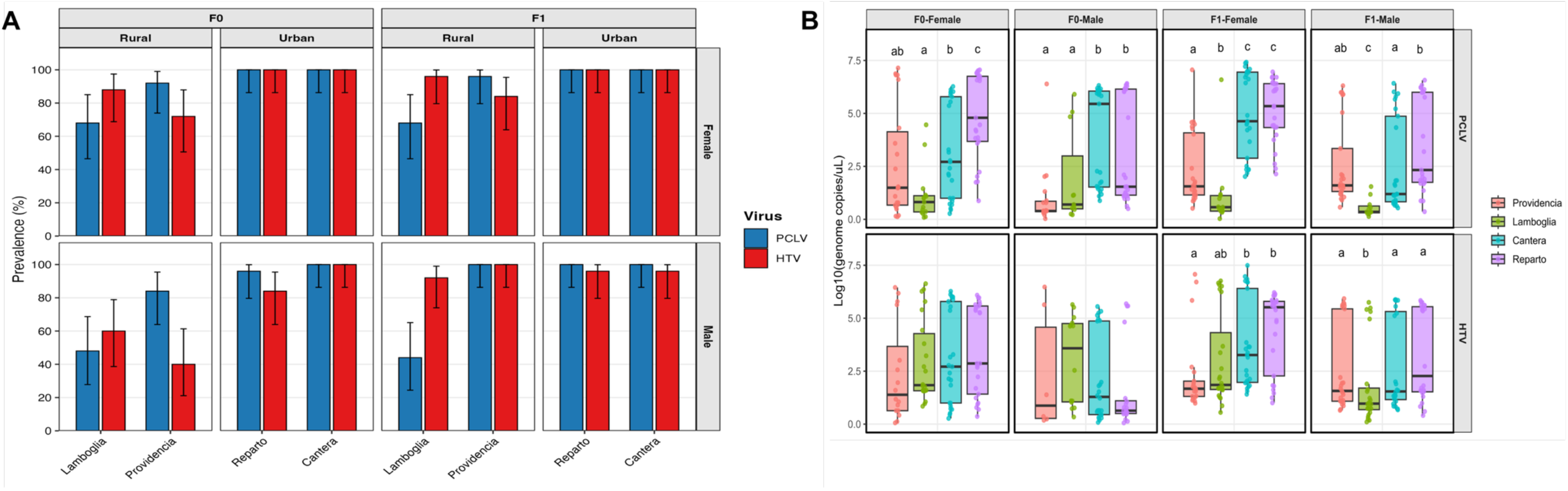
Prevalence and Viral genome copy numbers of Phasi Charoen-like virus (PCLV) and Humaita-Tubiacanga virus (HTV) across generations in urban and rural *Aedes aegypti* colonies. (**A)** Bars represent plots of the percentage prevalence of PCLV and HTV detected by RT-qPCR in F₀ and F₁ male and female mosquitoes from urban (Cantera and Reparto) and rural (Providencia and Lamboglia) colonies. F₀ and F₁ generations are shown separately to indicate the generation-level analysis structure. Each bar represents the percentage of virus-positive mosquitoes among 25 mosquitoes screened per Location × Generation × Sex × Virus group. Error bars indicate exact 95% binomial confidence intervals estimated using the Clopper–Pearson method implemented in R. **(B)** Distribution of PCLV and HTV genome copy numbers across *A. aegypti* populations in Puerto Rico. Boxplots show log₁₀-transformed genome copies/µL for four locations (Providencia, Lamboglia, Cantera, and Reparto), stratified by generation (F₀, F₁) and sex. Points represent individual mosquitoes, boxes indicate interquartile ranges, and horizontal lines represent medians. Statistical differences among locations within each panel were assessed using Kruskal–Wallis tests followed by pairwise Wilcoxon tests with BH correction. Panels with significant differences are annotated with compact letter displays; groups sharing letters are not significantly different.

GMV and ATLV showed distinct prevalence patterns between urban and rural mosquito populations (**Figure 7A**). Overall, GMV prevalence was consistently high in urban-derived colonies, with 100% detection in both females and males. In contrast, rural-derived colonies exhibited greater variability, with Lamboglia showing a higher infection prevalence (88–100%) while Providencia displayed markedly lower GMV prevalence (12– 16%), indicating heterogeneous circulation of GMV in rural settings. For ATLV, a clearer urban–rural contrast was observed. Urban colonies showed moderate prevalence, ranging from 56–80% in Reparto and 64–80% in Cantera across sexes. In comparison, rural colonies exhibited more variability and a lower infection prevalence, particularly in Providencia, where ATLV prevalence was reduced (28% in males and 68% in females). Although Lamboglia displayed high ATLV prevalence (72–100%), it remained more variable than the uniformly high GMV prevalence observed in urban samples.

**Figure 7.**
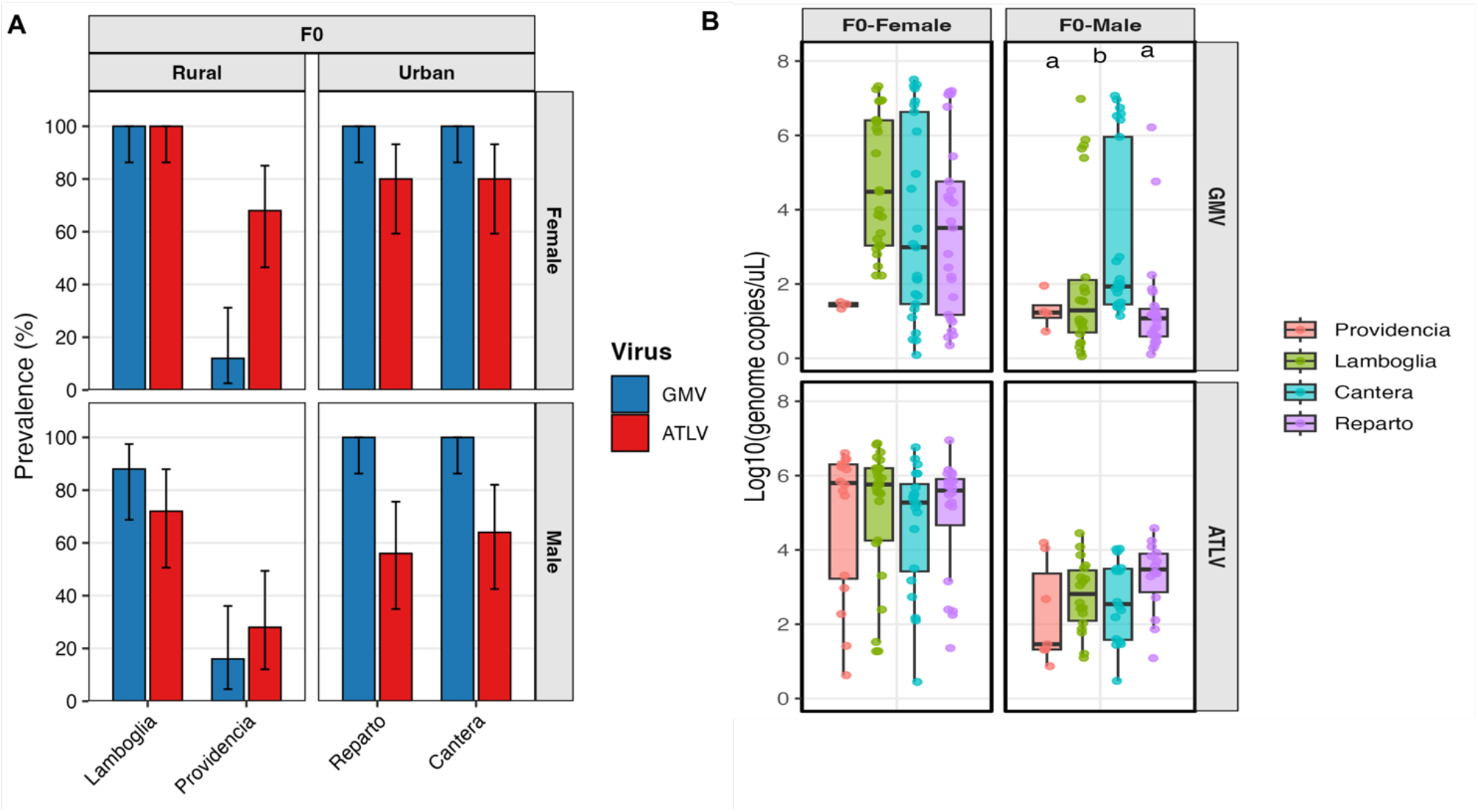
Prevalence and Viral genome copy numbers of Guadeloupe mosquito virus (GMV) and *Aedes aegypti* toti-like virus (ATLV) in urban and rural *Aedes aegypti* colonies. **(A)** Bars represent plots of the prevalence of GMV and HTV detected by RT-qPCR in F₀ male and female mosquitoes derived from urban (Cantera and Reparto) and rural (Providencia and Lamboglia). Prevalence was calculated as the proportion of positive individuals relative to the 25 mosquitoes tested per group. Error bars indicate exact 95% binomial confidence intervals estimated using the Clopper–Pearson method implemented in R. **(B)** Distribution of GMV and ATLV genome copy numbers across *A. aegypti* populations in Puerto Rico. Boxplots show log₁₀-transformed genome copies/µL for four locations (Providencia, Lamboglia, Cantera, and Reparto), stratified by generation (F₀) and sex. Points represent individual mosquitoes, boxes indicate interquartile ranges, and horizontal lines represent medians. Statistical differences among locations within each panel were assessed using Kruskal–Wallis tests followed by pairwise Wilcoxon tests with BH correction. Panels with significant differences are annotated with compact letter displays; groups sharing letters are not significantly different.

Consistent with ISV prevalence, virus loads were higher in urban than in rural mosquito populations **(Figure 6B).** For PCLV, urban populations (Cantera and Reparto) exhibited significantly higher genome copy numbers compared to rural populations across sexes and generations (Kruskal–Wallis, adjusted *p* < 0.05; **Figure 6B**). In contrast, HTV viral loads did not differ significantly between urban and rural populations across most comparisons (adjusted *p* > 0.05). Similarly, GMV and ATLV showed no significant differences in viral load between rural and urban mosquitoes (adjusted *p* > 0.05) **(Figure 7B).**

### Vector Competence Assay

*Infection rate:* Multilevel logistic regression revealed a significant effect of mosquito strain on infection status. We observed significantly higher odds of infection for the Reparto (urban strain) as compared to both Lamboglia (rural strain) and the laboratory strain ORL-D. Specifically, the odds of infection in Reparto were approximately 8.7-fold higher than in Lamboglia (OR = 8.70, 95% CI: 2.78–27.21, p = 0.0002). In contrast, no significant difference in infection odds was observed between ORL-D and Lamboglia (OR = 1.10, 95% CI: 0.52–2.31, p = 0.81), indicating comparable susceptibility between the laboratory and rural strains. Observed infection rates supported these model estimates, with Reparto showing consistently high infection across replicates (86.7–100%), compared to moderate infection rates in ORL-D (53.6–73.3%) and Lamboglia (60.0–63.3%) (**Figure 8**).

**Figure 8.**
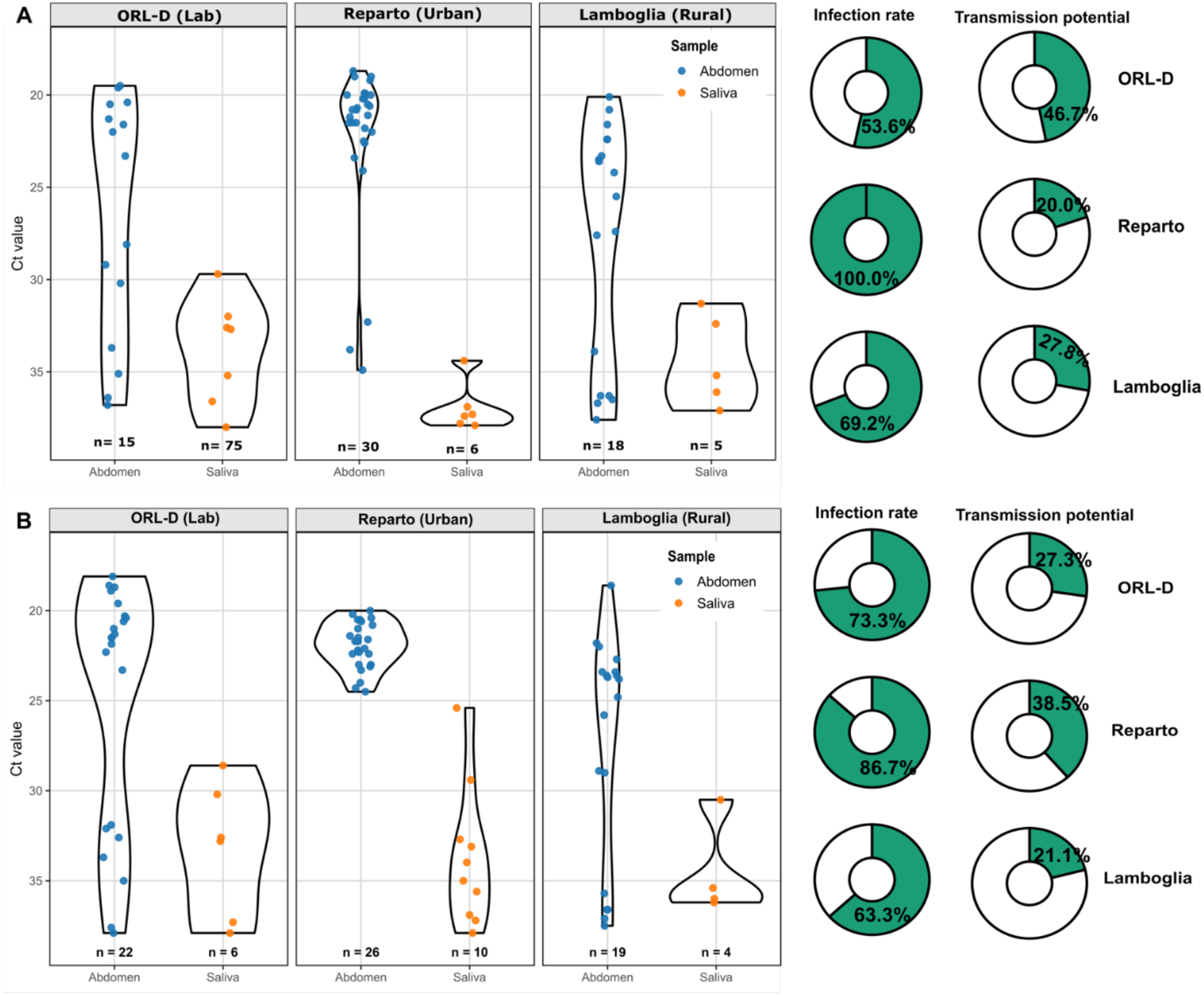
DENV-1 infection and transmission profiles across laboratory, urban, and rural *Aedes aegypti* strains. Violin plots show the distribution of RT-qPCR Ct values for DENV-1 in abdomen (blue; infection) and saliva (orange; transmission potential) samples from ORL-D (laboratory strain), Reparto (urban), and Lamboglia (rural) mosquitoes at 14 dpi. Data is presented for two experimental replicates A and B. Each point represents an individual mosquito, and violin shapes illustrate the distribution of Ct values within each group. Sample sizes (n) represent the number of positive samples for each tissue. Donut charts on the right summarize the proportion of positive samples for the abdomen (infection rate) and saliva (transmission potential) for each strain.

#### Transmission Potential

Unlike infection rates, mosquito strain did not significantly influence transmission potential. Although Reparto showed higher odds of saliva positivity compared to Lamboglia (OR = 1.90, 95% CI: 0.76–4.74, p = 0.17) and ORL-D (OR = 1.27, 95% CI: 0.29–2.17, Tukey-adjusted p > 0.05), these differences were not statistically significant. Similarly, no significant difference in transmission potential was detected between ORL-D and Lamboglia (OR = 1.51, 95% CI: 0.59–3.87, p = 0.39). Empirically, saliva positivity remained low across all groups, ranging from 13.3–33.3%, with the highest transmission rates observed in Reparto in replicate 2 (33.3%) (**Figure 8**).

### *Histone H4* expression analysis

It was recently shown that *histone H4* is a proviral host factor whose expression is maintained by the insect-specific viruses HTV and PCLV during DENV infection^4^. To determine if this host factor is influencing vector competence for Lamboglia and Reparto mosquitoes, we measured the mean expression levels for *histone H4* and observed an increased by 1.21-fold and 1.42-fold in Lamboglia and Reparto mosquitoes, respectively, relative to uninfected controls (**Supplemental Figure 1**). However, these differences were not statistically significant based on paired comparisons of ΔCt values following Benjamini– Hochberg correction (P > 0.05), indicating that DENV-1 infection did not significantly alter *histone H4* expression based on the resident ISV levels between the two colonies.

## DISCUSSION

Interest in the role of ISVs in the transmission of medically important viruses is driven by recent findings that natural co-infection of *A. aegypti* by both PCLV and HTV is associated with an increased (up to 200%) risk of dengue transmission^6^. In the past few years, several studies have investigated the distribution of ISVs *in Aedes* spp., given its broad geographical context and potential influence on arbovirus transmission risk^6,38–41^. Puerto Rico is a hotspot for dengue outbreaks, with typically differential disease incidence outcomes between urban and rural communities^19^. We conducted a microspatial analysis of ISV distribution among A. aegypti on Puerto Rico’s main island, sampling from both rural and urban settlements to investigate the impact of microscale diversity in A. aegypti viromes on vector competence for a regionally relevant DENV-1 strain.

### 1. ISV profiles of *A. aegypti* in Puerto Rico are geographically structured

Our metatranscriptomic analyses revealed clear differences in viral composition between urban and rural mosquito populations, with urban samples showing a higher proportion of viral reads compared to rural samples. This pattern is consistent with ecological studies demonstrating that mosquito viromes are shaped by habitat characteristics, environmental conditions, and host community structure ^39,42,43^. In a recent study across disturbance gradients, it was reported that changes in habitat composition drive turnover in mosquito host communities and their associated viral populations. The outcome of this ultimately influences virus prevalence patterns within vector species^42^. Similarly, a mosquito virome survey in the Philippines across study sites that differ in topography and land use transformations revealed that mosquito viromes are largely composed of ISVs, and that their diversity and abundance vary with land use, larval ecology, and environmental exposure^43^. Consistent with our findings, they observed greater diversity of viral families in sites that had transitioned from agricultural land to bare or built-up environments than in forested locations. This raises the important question of whether the differential abundance of ISVs in different ecological settings has an impact on mosquito vector competence for medically important arboviruses.

### 2. Urban *A. aegypti* have more stable core viromes

Our results suggest that PCLV, HTV, GMV, and ATLV are core components of the insect-specific virome of *A. aegypti* across two ecologically divergent sites in Puerto Rico. Among these, PCLV and HTV were the most consistently detected and predominant viruses across populations. Previous studies from Asia and South America have also reported a high abundance of PCLV and HTV in *A. aegypti* from dengue-endemic foci, suggesting that these viruses are commonly maintained in mosquito populations where dengue transmission occurs^6,43^. In contrast, co-circulation of these ISVs has been less frequently reported in African and European mosquito populations^6,9,42,44,45^. In our study, urban mosquitoes showed a more consistent, uniformly high prevalence of PCLV and HTV than rural mosquitoes. This pattern was especially evident for HTV, which showed lower and more variable prevalence in rural F₀ samples but increased prevalence in F₁ generations, suggesting stable vertical transmission over time. These observations indicate that these ISVs may be more stably established in urban colonies, while rural populations exhibit greater fluctuation in prevalence across sexes and generations^41,42^. Although clear differences in ISV prevalence and dominance were observed between urban and rural mosquitoes, it remains difficult to directly link these patterns to differences in vector competence or dengue incidence. For instance, although *A. aegypti* is abundant in Puerto Rico^45^ there are municipalities that do not report dengue cases even during epidemics^19^.

Both GMV and ATLV are the least characterized ISVs that were detected in the core virome of mosquitoes in Puerto Rico. In urban colonies, GMV prevalence was uniformly high across both sexes, whereas ATLV prevalence was consistently lower in males compared to females, particularly in Reparto (56% in males vs. 80% in females). A similar trend was observed in rural populations, where male prevalence was lower, especially in Providencia. Collectively, these results indicate that GMV is more stably maintained in urban mosquito populations, with near fixation across colonies, whereas rural populations show greater variability, particularly in Providencia, where low prevalence drives this variability. Considering that GMV was detected in humans with febrile illness^10^ and its high prevalence in *A. aegypti,* clinical surveillance efforts should monitor for GMV among febrile patients who are negative for other endemic arboviruses. In contrast, ATLV demonstrates moderate prevalence in urban colonies but a more heterogeneous, generally reduced prevalence in rural populations, suggesting differential maintenance dynamics of this ISV between urban and rural mosquito populations. ATLV has also been identified in field strains in Florida, USA^15^ and laboratory strains^46^ of *A. aegypti*; however, its influence on vector competence remains to be explored.

We detected CFAV in all field and laboratory mosquito colonies at very low abundance (<1%), except in ORL-D, where CFAV was the only ISV identified. Historically, CFAV was the dominant ISV in Puerto Rico in the late 1990’s^20^; however, our data suggest a transient reduction of CFAV prevalence and a shift toward PCLV and HTV as the dominant ISVs. Given that CFAV has been reported to antagonize DENV dissemination in mosquitoes^22^ this shift in the ISV profile over several decades may have contributed to the transmission of multiple DENV serotypes during this period^6,48^.

### 3. The effect of PCLV–HTV co-infection on DENV-1 transmission is inconclusive

Vector competence studies revealed that the urban Reparto strain had significantly higher odds of DENV-1 infection than the rural Lamboglia strain and the laboratory ORL-D strain. However, saliva positivity did not differ significantly across strains, indicating that infection and transmission phenotypes were decoupled. Our findings do not align with prior studies, which found that PCLV and HTV were associated with enhanced arbovirus transmission^6^. This disparity could be due to several factors, including differences in host genetic background, microbiome composition, and the DENV serotype used^3,47,48^. For instance, a recent report shows that PCLV reads were significantly different between DENV-1- and DENV-2-infected mosquitoes, suggesting within-host modulation of ISVs during the course of infection^48^. Furthermore, arbovirus transmission requires successful progression through multiple barriers, including midgut infection, dissemination, and salivary gland infection and escape^5,49^. Differences in infection rates without corresponding differences in transmission potential suggest that mosquito virome-associated factors may primarily influence early infection processes rather than later transmission stages^49^. Emerging literature indicates that ISVs can modulate arbovirus replication within mosquito hosts through mechanisms such as immune priming, resource competition, or interference with viral replication pathways^5–7,11^. While our study was not designed to experimentally manipulate ISVs, the observed association between distinct ISV profiles and increased infection susceptibility in urban mosquitoes is consistent with the hypothesis that virome composition may contribute to variation in vector competence. However, causality cannot be inferred from the present data.

#### DENV-1 infection resulted in modest changes in *histone H4* expression in both mosquito colonies

Given that *histone H4* has been identified as a proviral host factor whose expression is modulated by PCLV and HTV^6^, we investigated whether differences in ISV composition between the Lamboglia and Reparto colonies were associated with altered *histone H4* expression following DENV-1 infection. *Histone H4* expression was modestly elevated in both colonies, with a greater increase observed in Reparto than in Lamboglia mosquitoes. Although these differences were not statistically significant, the trend is consistent with Olmo et al., who showed that HTV and PCLV prevent DENV-mediated downregulation of *histone H4* and that *histone H4* facilitates DENV replication. Since Reparto mosquitoes harbored higher prevalences of both HTV and PCLV than Lamboglia, the higher fold-change observed in Reparto may reflect stronger ISV-associated modulation of host transcriptional responses. However, the small magnitude of change and lack of statistical significance suggest that this effect was subtle under the conditions tested. Direct comparison with Olmo et al. is limited because our analysis compared colonies differing in HTV/PCLV prevalence rather than individual HTV/PCLV-positive and -negative mosquitoes. In addition, Olmo et al. observed significant *histone H4* modulation at 8 dpi but not 14 dpi. We tested and analyzed at 14 dpi since this time point is associated with maximal salivary gland infections. However, due to sampling differences, any ISV-associated effect may have occurred earlier and no longer be detectable at the time of sampling.

### Limitations

Although the results support a link between ISV composition and infection susceptibility, several alternative explanations must be considered. Differences in host genetic background and microbiome composition between urban and rural colonies could independently influence vector competence^5,50,51^. Additionally, metatranscriptomic analyses reflect viral RNA abundance rather than active replication alone and may be influenced by sequencing depth, host RNA depletion efficiency, and database annotation biases. Furthermore, we used DENV-1 (2014, genotype V) for the vector competence assay; however, recent data indicate it may not have been the predominant serotype in Puerto Rico during the sampling period^52^.

### Conclusions

In sum, our findings support the broader ecological framework that mosquito viromes are shaped by environmental context and that these ecological differences may influence *A. aegypti* vector competence for dengue viruses at smaller spatial scales than previously appreciated. Habitat-driven variation in ISV composition, particularly shifts in dominant ISVs, could contribute to spatial heterogeneity in dengue transmission risk within endemic regions. Importantly, our data suggest that higher infection rates in urban mosquito populations in Puerto Rico may be associated with stable, high-prevalence ISV communities. This observation highlights the need to consider the mosquito virome as an integral ecological component of vector competence rather than a passive background microbiome. To establish a causal role for ISVs in shaping vector competence, future studies should incorporate experimental manipulations of the mosquito virome, including ISV clearance, virome transplantation, and complete adaptation, prior to controlled co-infection experiments. Our analysis supports our prevailing hypothesis that geographical partitioning influences ISV diversity at the community level, compelling future functional studies to evaluate the effects of varying ISV profiles on arbovirus transmission by geographically segregated mosquito populations worldwide.

## Data Availability

Raw sequencing data have been deposited in the NCBI Sequence Read Archive under BioProject accession number **PRJNA1054025**. Assembled viral genome sequences have been deposited in GenBank under accession numbers **PZ595439–PZ595477**, **PZ669725– PZ669763**, and **PZ669925–PZ669934**.

## Code Availability

No new custom code was generated for this study.

## Acknowledgements

This research was supported by the Southeastern Regional Center of Excellence in Vector-borne Diseases: The Gateway Program, which is funded by the cooperative Agreement U01CK000662 from the Centers for Disease Control and Prevention. The contents are solely the responsibility of the authors and do not necessarily represent the official views of the Centers for Disease Control and Prevention.

## Author contributions

Conceptualization: B.A., R.R.D. Methodology: B.A., R.R.D.

Sampling: N.N.-M., J.M.Q., N.B.-S., V.R.-A., R.R.-G., G.B., R.R.D.

Mosquito colony maintenance and infection experiments: B.A., L.A.P, J.F.W.

Molecular assays and data generation: B.A.

Bioinformatic and statistical analyses: B.A., H.J.B.

Data visualization: B.A.

Writing—original draft preparation: B.A.

Writing—review and editing: B.A., L.A.-P., H.J.B., N.N.-M., J.M.Q., N.B.-S., J.F.W, V.R.-A., R.R.-G., G.B., R.R.D.,

Supervision and Funding acquisition: R.R.D.

