## Supplementary Table 2 for "Microspatial partitioning of insect-specific viromes and dengue virus transmission risk by *Aedes aegypt*i in Puerto Rico"

| Supplementary Table 2 – List of oligonucleotides used in this study. | | |
|---|---|---|
| **Target** | **Description** | **Sequence (5'-3')** |
| Phasi Charoen-like phasivirus | PCLV-RdRp FWD | GAACTGGAGAGCCCAGTTGATT |
| | PCLV-RdRp REV | CAGCTCCTTCAACGAAAGTGAC |
| | PCLV-RdRp Probe | TACACTAGCTAGCTCACTGCCTGGAG |
| Humaita-Tubiacanga virus | HTV-Capsid FWD | TGGTCAAGATGGCGGCCTCGAG |
| | HTV-Capsid REV | AAGTCTGCGGAGGTGTTATTTGC |
| | HTV-Capsid Probe | ACTTATGGGTCGTTGGTTGTGGGTTG |
| Guadeloupe Mosquito Virus | GMV-Seg-2_ORF2 FWD | ACGTCTCTGTTGCACTTTAGGG |
| | GMV-Seg-2_ORF2 REV | ACTGTCGAGCACTTAACGACAC |
| | GMV-Seg-2_ORF2 Probe | AGTTCCGCAGCGAACAGTAGCGAGTCG |
| *Aedes aegypti* toti-like virus | ATLV-RdRp FWD | CATCGCCCTCAATAGCCAAATC |
| | ATLV-RdRp REV | CGTCATAGGCGGTCCTAGATCC |
| | ATLV-RdRp Probe | CCATAACCATTCCGGTGGCATACCTC |
| *Ae. aegypti Histone H4* | Histone H4 FWD | CTACCGTCGTCGAACACAGA |
| | Histone H4 REV | CCTCCTTTTCCGAGTCCTTT |
| *Ae. aegypti rpl32* | rpl32 FWD | CAGTCCGATCGCTATGACAA |
| | rpl32 REV | ATCATCAGCACCTCCAGCTC |
| Dengue Virus 1 | DENV-1 FWD | CAAAAGGAAGTCGYGCAATA |
| | DENV-1 REV | CTGAGTGAATTCTCTCTGCTRAAC |
