## Supplementary Table 1 for "Microspatial partitioning of insect-specific viromes and dengue virus transmission risk by *Aedes aegypt*i in Puerto Rico"

| Supplementary Table 1 – Read mapping statistics of analyzed samples. | | | | | | | | | | | | |
|---|---|---|---|---|---|---|---|---|---|---|---|---|
| **Sample** | **Designation** | **Raw reads (N)** | **Filtered non-host reads** | | **Total non-host Reads aligned to DB** | | **Non-host Reads aligned to DB** | | | | | |
| | | | | | | | **Eukaryotic** | | **Bacterial** | | **Viral** | |
| | | | **Number (N)** | **Percentage (%)** | **Number (N)** | **Percentage (%)** | **Number (N)** | **Percentage (%)** | **Number (N)** | **Percentage (%)** | **Number (N)** | **Percentage (%)** |
| CMAVE | Lab | 43,729,370 | 9,613,067 | 21.98 | 262,579 | 2.73 | 27,864 | 10.61 | 122,019 | 46.47 | 101,977 | 38.84 |
| ORL-D.1 | Lab | 66,640,717 | 4,981,208 | 7.47 | 159,102 | 3.19 | 110,930 | 69.72 | 40,551 | 25.49 | 94 | 0.06 |
| L4 | Rural | 43,200,614 | 2,769,847 | 6.41 | 240,641 | 8.69 | 32,217 | 13.39 | 182,268 | 75.75 | 30,417 | 12.64 |
| P1 | Rural | 41,933,024 | 3,108,533 | 7.41 | 937,149 | 30.15 | 37,576 | 4.01 | 52,703 | 5.62 | 360,821 | 38.5 |
| P3 | Rural | 47,175,041 | 3,052,903 | 6.47 | 308,382 | 10.1 | 26,604 | 8.63 | 157,596 | 51.1 | 123,208 | 39.95 |
| C4 | Urban | 40,221,924 | 3,404,292 | 8.46 | 1,146,183 | 33.67 | 73,894 | 6.45 | 5,336 | 0.47 | 1,059,109 | 92.4 |
| R2 | Urban | 43,793,709 | 4,783,356 | 10.92 | 2,391,919 | 50.01 | 33,924 | 1.42 | 37,575 | 1.57 | 1,761,097 | 73.62 |
| R4 | Urban | 40,295,636 | 4,606,650 | 11.43 | 2,529,209 | 54.9 | 26,014 | 1.03 | 164,816 | 6.52 | 2,334,457 | 92.3 |
