## Supplementary Figure 1 for "Microspatial partitioning of insect-specific viromes and dengue virus transmission risk by *Aedes aegypt*i in Puerto Rico"

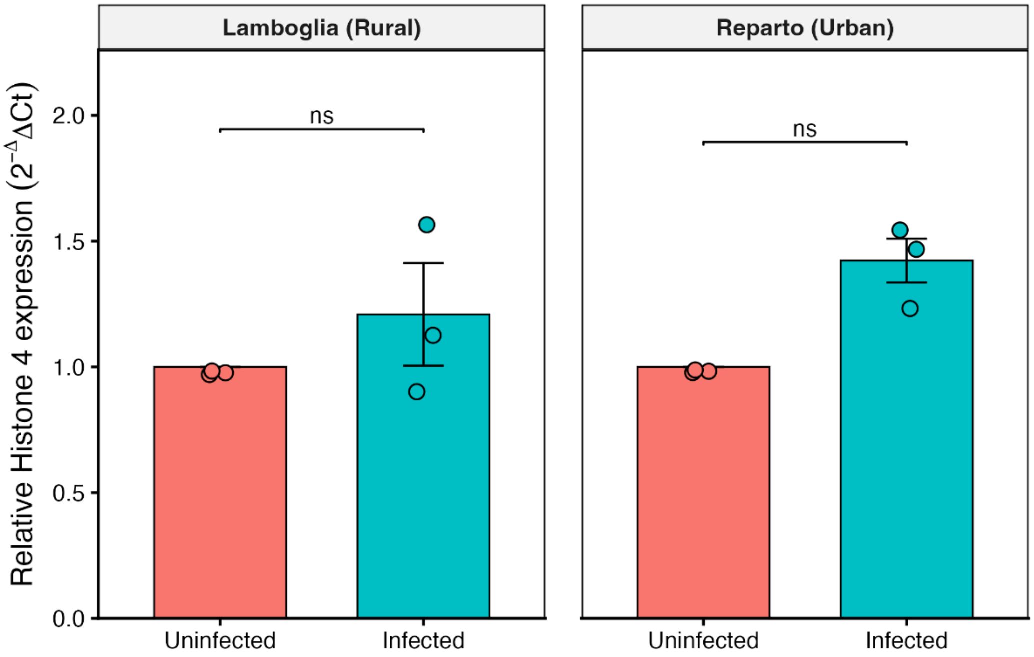


**Supplementary Figure 1. Relative *Histone H4* expression following DENV-1 infection in Lamboglia and Reparto mosquito colonies.** Relative expression levels were determined by RT-qPCR using the 2^-ΔΔCt method with uninfected samples as the reference group. Bars represent mean expression values ± SEM, while points indicate individual biological replicates (n = 3). Statistical comparisons between infected and uninfected groups were performed using paired Student's t-tests on ΔCt values with Benjamini–Hochberg correction for multiple testing. All statistical analyses and data visualizations were performed in R using the ggplot2, ggpubr, and rstatix packages.
